# Generation of dual specific bivalent BiTEs (dbBIspecific T-cell Engaging antibodies) for cellular immunotherapy

**DOI:** 10.1101/559427

**Authors:** Maciej Kujawski, Lin Li, Supriyo Bhattacharya, Patty Wong, Wen-Hui Lee, Lindsay Williams, Harry Li, Junie Chea, Kofi Poku, Nicole Bowles, Nagarajan Vaidehi, Paul Yazaki, John E. Shively

## Abstract

Bispecific T-cell engaging antibodies (BiTES), comprising dual anti-CD3 and anti-tumor antigen scFv fragments, are important therapeutic agents for the treatment of cancer. The dual scFv construct for BiTES requires proper protein folding while their small molecular size leads to rapid kidney clearance. Here we show that an intact (150 kDa) anti-tumor antigen antibody to CEA was joined in high yield (ca. 30%) to intact (150 kDa) anti-murine and anti-human CD3 antibodies using hinge region specific Click chemistry to form dual-specific, bivalent BiTES (db BiTES, 300 kDa). The interlocked hinge regions are compatible with a structural model that fits the electron micrographs of the 300 kDa particles. Compared to intact anti-CEA antibody, dbBiTES maintain high in vivo tumor targeting as demonstrated by PET imaging, and redirect dbBiTE coated T-cells (1 microgram/10 million cells) to kill CEA^+^ target cells both in vitro, and in vivo in CEA transgenic mice.

## Introduction

The concept of directing T-cells to tumors with bispecific antibodies that incorporate an anti-CD3 antibody and an anti-tumor antigen antibody into a single molecule was first explored 25 years ago when recombinant DNA technologies made the genetic engineering of bispecific antibodies possible [1]. Today the most widely used BiTE (Bispecific T-cell Engaging antibodies) constructs join two single chain Fv fragments together with a linker to form a 50kda protein. Examples of tumor target antigens include CD19, EpCAM, Her2/neu, EGFR, CEA, and more [1]. The BiTE anti-CD19/anti-CD3 (Blinatumomab) has had remarkable success in the treatment of several B-cell malignancies, suggesting that directing T-cell (and other effector cells) to tumors by a bispecific antibody approach can be effective even against solid tumors [2]. Possible limitations of the therapy include the small molecular size that requires constant infusion of rather large amounts of BiTEs, the poly-activation of endogenous TCRs may lead to off target toxicity, and the lack of an engagement of a co-stimulus such as CD28 on the T-cell may limit effective tumor killing. An alternate approach pioneered by Lum and associates [3] utilized chemical conjugation with protein surface cross-linking agents on intact antibodies (eg., anti-CD3 and anti-CA125) that were coated onto IL-2 activated autologous T-cells. In spite of the high heterogeneity of the resulting bispecific antibodies, advantages of the approach were that only microgram amounts of antibody were required for coating the T-cells and the therapy was considered a cell-based therapy by the FDA since it involved infusion of T-cells and not antibody infusion. Thus, this approach used clinically tested humanized antibodies as a starting point, avoiding the need to engineer dual scFVs that may be difficult to express in the amounts needed for therapy. Furthermore, the infusion of bispecific antibody coated T-cells is a cell-based therapy similar, and perhaps complimentary, to infusion of CAR T cells. However, to bring the approach into widespread use, it is important to develop a high yield conjugation method that does not lead to a complex mixture of bispecific antibodies.

To overcome this challenge, we show here an efficient approach to conjugate two antibodies alkylated at their sulfhydryl reduced hinge regions with two complimentary Click reagents, bromoacetamido-dibenzocyclooctyne (DBCO) and bromoacetamido-PEG_5_-azide. To demonstrate feasibility, we selected the humanized anti-CEA antibody T84.66-M5A (M5A) and anti-CD3 antibody OKT3, both extensively used in the clinic [4, 5]. Given their **d**ual specifities and retention of **b**ivalent binding, we propose to call them dbBiTEs. The 300 kDa product when purified away from contaminating 150 kDa species, shows distinctive six-lobed antibody particles on electron microscopy that fit atomic scale models, labels both CEA and CD3 positive cells, and when coated on activated T-cells, kills CEA positive cells in cytotoxicity assays. In vivo PET imaging shows excellent targeting to tumor targets, and in the preliminary therapy studies, killing of CEA positive tumors.

## Results

### Generation of dbBiTES

Building on the approach developed by Lum et al. [3] who randomly cross-linked two intact antibodies (anti-CD3 and anti-CA125) using Traut’s reagent and sulfosuccinimidyl 4-(*N*-maleimidomethyl) cyclohexane-1-carboxylate to redirect T-cell therapy to tumor targets, we first generated bispecific antibodies by cross-linking two intact antibodies at their hinge regions using Click chemistry (**Scheme 1**). The use of Click chemistry allows the generation of a bispecific antibody product with improved yields and molecular characteristics, a much-desired feature in bringing new products to the clinic. To distinguish the product from conventional BiTEs built from monovalent single chain Fv (scFv) fragments, we suggest the name **dbBiTE**, for **d**ual specific **b**ivalent **BiTE.** Briefly, OKT3, a murine anti-human CD3 antibody widely used in the clinic [4] was reduced at its hinge region cysteines under non-denaturing conditions, alkylated at its reduced hinge region cysteines with a bromoacetamido-PEG_5_-azido derivative and conjugated to our humanized anti-CEA M5A antibody [5] that was similarly reduced and alkylated with a bromoacetamido-DIBO derivative. Each of the derivatized IgGs were analyzed by electrospray ionization mass spectrometry to confirm their degree of derivatization (**Supplementary Fig S1**). In both cases the heavy chains contained at least 2 Click derivatives. The Clicked dbBiTE was isolated in a yield of 30% by size exclusion chromatography (SEC, peak 1, 300 kDa, **Fig 1A**). A second peak of MW 150 kDa was obtained in a yield of 60%. The peaks were analyzed by non-denaturing SDS gel electrophoresis to further verify their molecular sizes compared to the two intact antibodies (**Fig 1B and C**). The 150 kDa peak was shown to be mainly a monovalent bispecific antibody that was poorly active in functional studies (redirected T-cell cytotoxicity) and was broken down into lower molecular weight fragments by SDS gel electrophoresis (data not shown). Therefore, we focused on the novel dbBiTE (300 kDa) since it retains the inherent avidity (bivalent binding) of both parent antibodies. Peak 1 was re-purified by SEC to remove contaminating peak 1. Rechromatography of peak 1 over the course of several weeks showed no evidence of instability consistent with their covalent linkages (data not shown).

**Scheme 1.** Cysteine hinge specific Click chemistry for generation of dbBiTES. Blue = reduced antibody 1 with DBCO. Red = reduced antibody 2 with PEG_n_-azide. Click = the two derivatized antibodies were mixed 1:1. The cross-linking between two heavy chains is likely the favored result, including the possibility of two heavy chain cross-links per dbBiTE. Evidence for cross-linking between two light chains (side to side) is shown in the EM studies as a rare event.

**Figure 1.**
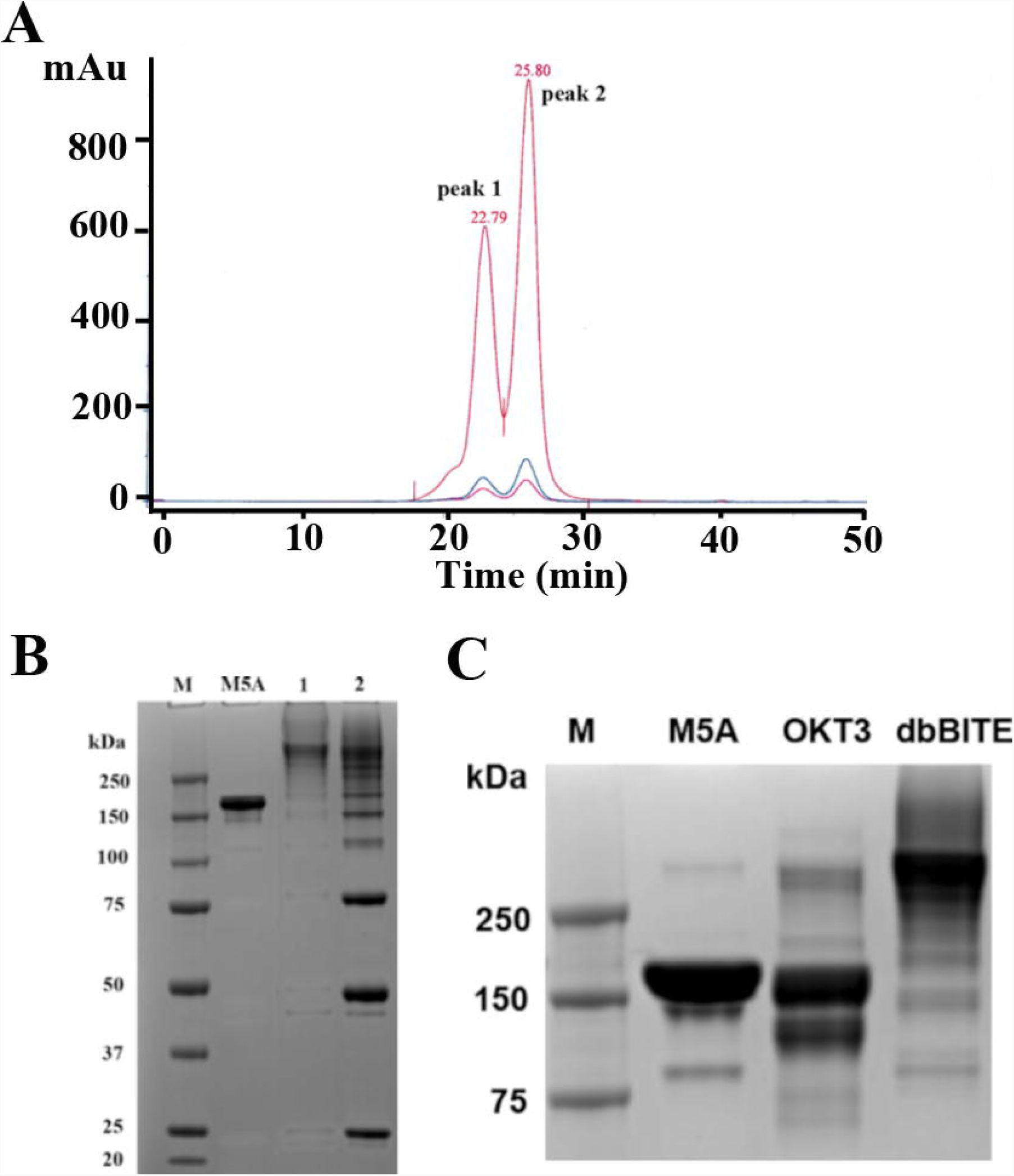
Purification of a dbBiTE by size exclusion HPLC. **A**. One hundred micrograms of a crude reaction mixture of DBCO-anti-CD3 antibody plus azido-PEG5-anti-CEA antibody were injected onto a Superdex 200 (1 × 30 cm) column, monitored at 214 nm (red) and 280 nm (blue) and eluted at a flow rate of 0.5 mL/min in PBS. The two peaks (peak 1 and 2) were collected and further analyzed. Based on calibration with authentic standards, peak 1 has a molecular mass of 300 kDa and peak 2, 150 kDa. **B**. Peaks 1 and 2 from SE HPLC were run on non-reducing SDS gels to determine their molecular sizes compared to the standard IgG M5A. **C**. The starting antibodies, anti-CEA M5A and anti-CD3 OKT3 were run as standards on SDS polyacrylamide gel electrophoresis alongside purified peak 1 dbBiTE.

### Particle size by electron microscopy

Purified peak 1 was analyzed by transmission electron microscopy (TEM) to determine the particle size and morphology (**Fig 2A**). Most particles exhibited a 5-lobed morphology suggesting that the predicted sixth lobe (based on 3 lobes per Ab times 2) was hidden under or above the plane of the 2D image. This morphology is more clearly visualized on the 2D averaged analysis (**Fig 2B**). Using tilt axis imaging, it was possible to show a representative 3D image (**Fig 2C**). A closer examination of the particles shown in **Fig 2A** reveals multiple orientations of 6-lobed particles (further data available upon request), always with the sixth lobe appearing as additional density in the center of the particle. These images are compatible with the random landing of 6-lobed 3D particles on a 2D grid in which one lobe is lying above or below the main body of the particle. In addition to the majority of 6-lobed particles found, a small percentage (<10%) of particles with a side-to side orientation were observed. We speculate that these are due to Click chemistry occurring between adjacent light chains (1 cysteine per light chain) at their hinge regions, rather than the more frequent heavy chain hinge regions (3 cysteines per heavy chain in human IgG1).

**Figure 2.**
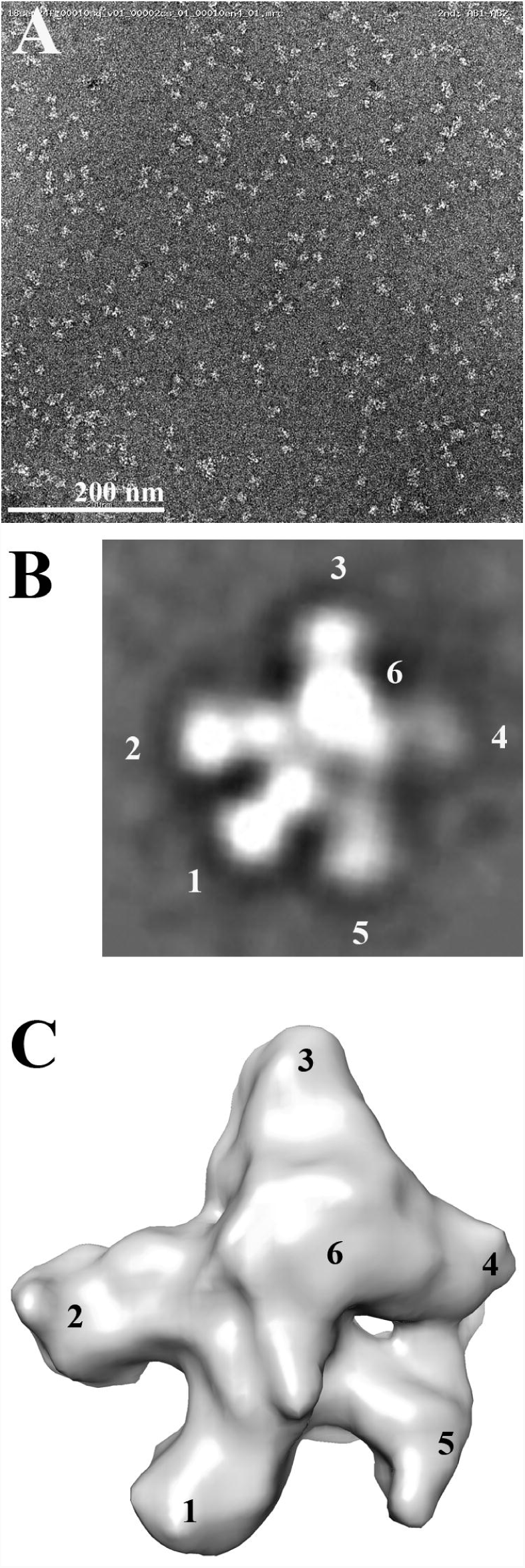
Electron microscopy analysis of dbBiTES. **A**. Representative EM of 300 kDa particles at 67,000 magnification. **B**. 2D average reconstruction of >100 6-lobed particles. IgG lobes arbitrarily labeled as 1-6, the 6^th^ lobe assumed out of the plane (z-axis) of the paper. **C**. 3D reconstruction of 6-lobed particles using tilt beam EM. Lobes labeled as above.

### Molecular simulations of dbBiTE structure

We performed multi-scale molecular dynamics simulations to generate an atomic level structural model examining how the two IgGs fit into a dbBiTE joined by Click chemistry at their hinge region cysteines. We generated a coarse grain homology model of the two IgGs (**Supplemental Fig S2)** that used the coarse grain simulation method *Martini* to optimize the packing of these two moieties. The dynamics of the two IgGs approaching each other and docking at their hinge regions is shown in **Supplemental movie SM1**. The optimized structural model shows the two IgGs joined by at least two pairs of Clicked hinge region cysteines (**Fig 3 and supplemental movies SM2 and 3**). The final docked dbBiTES were then oriented to fit within the observed 6-lobed particles found on EM (**Supplemental Fig S3**). The superposition of the dbBiTE structural model thus generated, onto the particles imaged by EM strongly supports the idea that two IgGs can indeed be joined together at their hinge regions in spite of the size of their 3 globular domains.

**Figure 3.**
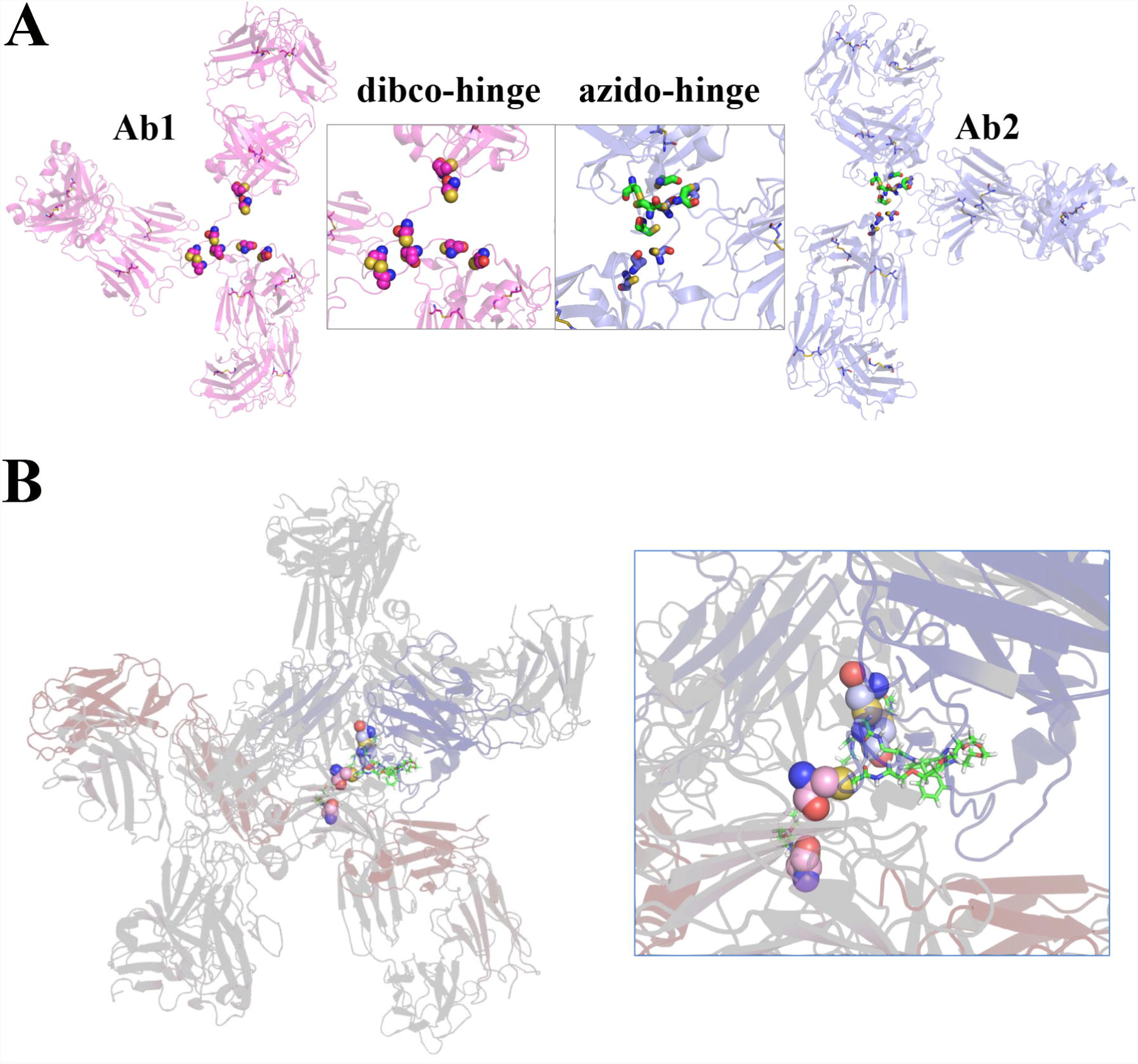
Structural model of the dbBiTE. **A**. Details of the hinge region. The two Clicked reagents, DBCO and PEG_5_-azide are shown attached to cysteines in the hinge regions of two IgG1s. **B**. Structural model derived after coarse grain MD simulations showing possible Clicked derivatives between two pairs of cysteines in the heavy chain hinge of a dbBiTE.

### In vitro binding and cytotoxicity of dbBiTEs to CEA positive targets

Since it was important to demonstrate that both antibody specificities were retained in the dbBiTE, in vitro binding studies were performed comparing the starting antibodies to the dbBiTE on CEA and CD3 positive targets (**Fig 4A-B**). The results demonstrate that dbBiTES are able to bind both CEA and CD3 positive target cells. In vitro cytotoxicity was demonstrated by coating activated human T-cells with dbBiTES (1 μg per 10M cells per mL) and incubation with CEA positive targets at the indicated E:T ratios (**Fig 4C**). Effective killing was observed as low as an E:T of 1.25:1 with maximal killing at an E:T of 10:1. Analysis of the supernatants revealed a significant release of IFNγ compared to controls (**Fig 4D**) demonstrating that the dbBiTE coated activated T-cells were able to produce a functional cytokine in response to target engagement. When the coating capacity of activated T-cells with dbBiTEs was tested by flow analysis, it was found that as little as 1 ng/mL of dbBiTE incubated with 10M T-cells per mL was detectable (**Fig 4E**). Although the cytotoxicity of activated T-cells against CEA positive targets was detectable at this concentration, higher coating concentrations were more effective (**Fig 4F**). Microscopic images of the killing of CEA positive targets by dbBiTE coated activated T-cells are shown in **Supplementary Fig S4** and **Supplementary movies SM4 and 5**.

**Figure 4.**
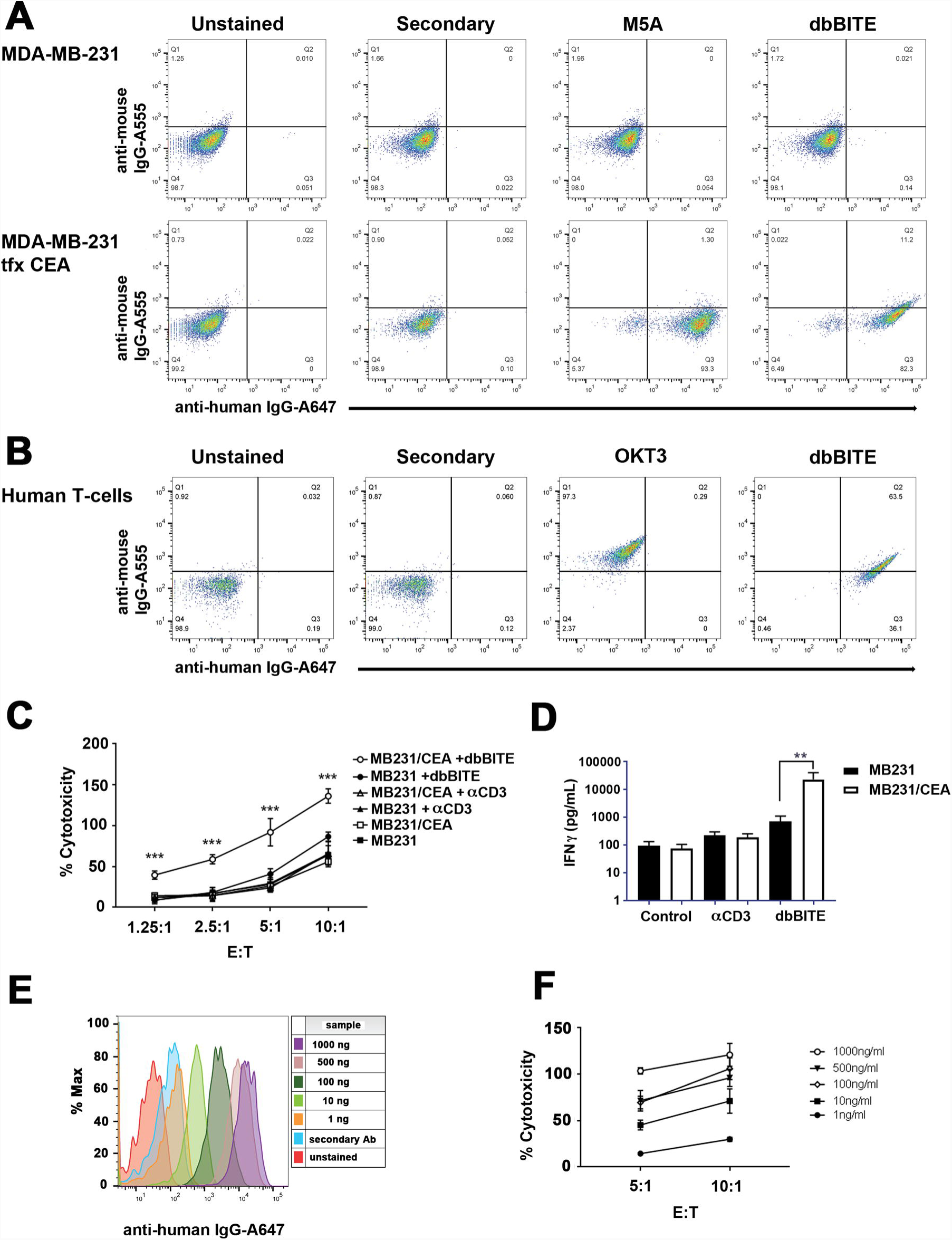
In vitro binding of dbBiTEs to CEA positive target cells and to human T-cells. **A.** Flow analysis of anti-CEA antibody M5A and dbBiTE to CEA positive MDA-MB-231 cells. **B.** Flow analysis of anti-CD3 antibody OKT3 and dbBiTE to CD3 positive human T-cells. **C**. Activated human T-cells coated with dbBiTE were incubated with target cells that expressed CEA (MB231CEA) or not (MB231) at indicated E:T ratios and cytotoxicity measured by an LDH release assay (details in Supplement). Controls are uncoated or anti-CD3 antibody coated T-cells. The results show that only dbBiTE coated T-cells exhibit significant cytotoxity towards CEA+ and not CEA-targets, p<0.001. Antibody concentrations were 1 μg/mL per ten million cells. **D**. IFNγ secretion was measured in the media at 24 hrs at an E:T ratio of 10:1. Controls include uncoated T-cells or T-cells coated with anti-CD3. The assay was performed in triplicate, p<0.01. **E**. Binding of dbBiTE to human T-cells was measured at concentrations ranging from 1 μg/mL to 1 ng/mL. **F**. Cytoxicity was measured for T-cells coated at 1 μg/mL to 1 ng/mL against MDA-MB231-CEA target cells at two E:T ratios.

### In vivo targeting of dbBiTEs to a CEA positive tumor in CEA Tg mice

It was also important to demonstrate in vivo targeting of dbBiTEs to CEA positive tumors, since therapy studies were planned with dbBiTE coated T-cells. This study involved DOTA conjugated dbBiTEs labeled with ^64^Cu for positron emission tomography (PET) of CEA positive human colon carcinoma LS174T xenografts in NOD/SCID mice. When radiolabeled dbBiTE (300 kDa) was compared to intact M5A (150 kDa) on SEC, dbBiTE was shown to be about 88% single species with about 12% molar contamination of a 150 kDa species (**Supplementary Fig S5**). The 150 kDa contaminant is likely a mixture of intact M5A and OKT3 or their Click conjugated half molecules. Since only the 300 kDa species shifted to a higher molecular complex on the addition of unlabeled CEA, we conclude that only the 300 kDa dbBiTE is immunoreactive.

When ^64^Cu-DOTA labeled dbBiTE was injected into CEA positive tumor bearing mice, PET imaging revealed rapid uptake into both tumor and liver with evidence of blood pool in the heart at the earliest time point (**Supplementary Fig S6**). At the terminal time of 44 hr, 15% ID/g was found in tumor, 5% ID/g in blood and 18% ID/g in liver. Compared to our previously published PET images of ^64^Cu-DOTA labeled M5A that showed about 45% ID/g in tumor, 10% ID/ in blood and 12% ID/g in liver at 44 hrs [6], the faster clearance of dbBiTE from blood into liver may be due to liver recognition of the multiple Fc moieties on the dbBiTE. Nonetheless, the PET imaging demonstrates excellent targeting of the dbBiTE to tumor, encouraging us to test the efficacy of the dbBiTE in vivo.

### In vivo killing of CEA positive tumors

A limited therapy study of dbBiTE coated T-cells was performed in CEA transgenic mice bearing the syngeneic colon cancer cell line MC38 transfected with CEA and luciferase. This model (minus the luciferase) has been previously described by us using the immunocytokine anti-CEA-IL-2 [7]. In order to test a biocompatible version of the dbBiTE in this model system, we generated a dbBiTE comprising rat anti-mouse CD3 and anti-CEA M5A (data not shown).

The in vitro cytotoxicity of this dbBiTE was first demonstrated with two CEA transfected cell lines, murine mammary carcinoma line E0177 and murine colon carcinoma line MC38, compared to untransfected parental cells (**Supplementary Fig S7**). Tumors were inoculated i.p. and 3 mice were selected as controls (little or no luciferase expression after 9 days) and 3 with carcinomatosis (**Fig 5A**). The mice were treated i.p. four times every three days with 10M activated murine spleen T-cells either coated or not with 1 μg of dbBiTE. Four days after the last injection of dbBiTE coated T-cells, the mice were euthanized and examined for the presence of tumor nodules. The number of tumor nodules that were found either peri-pancreatic or peri-intestinal are shown in **Fig 4B** along with micrographs shown in **Fig 5C** and **E**. Tumor nodules were then digested and analyzed by flow for the presence of CEA, CD45 lymphocytes, CD4 or CD8 T-cells, and Ly6G neutrophils or F4-80 macrophages (**Fig 4D**). The results demonstrate an absence of CEA in dbBiTE treated tumors and infiltration of Ly6G^+^ neutrophils, while CEA is present in controls along with infiltration of F4/80^+^ macrophages. Although this is a very preliminary therapy study, we are encouraged that dbBiTE coated T-cells may have therapeutic potential, and with further development, become an alternative to the genetic manipulation necessary with CAR T-cell therapy.

**Figure 5.**
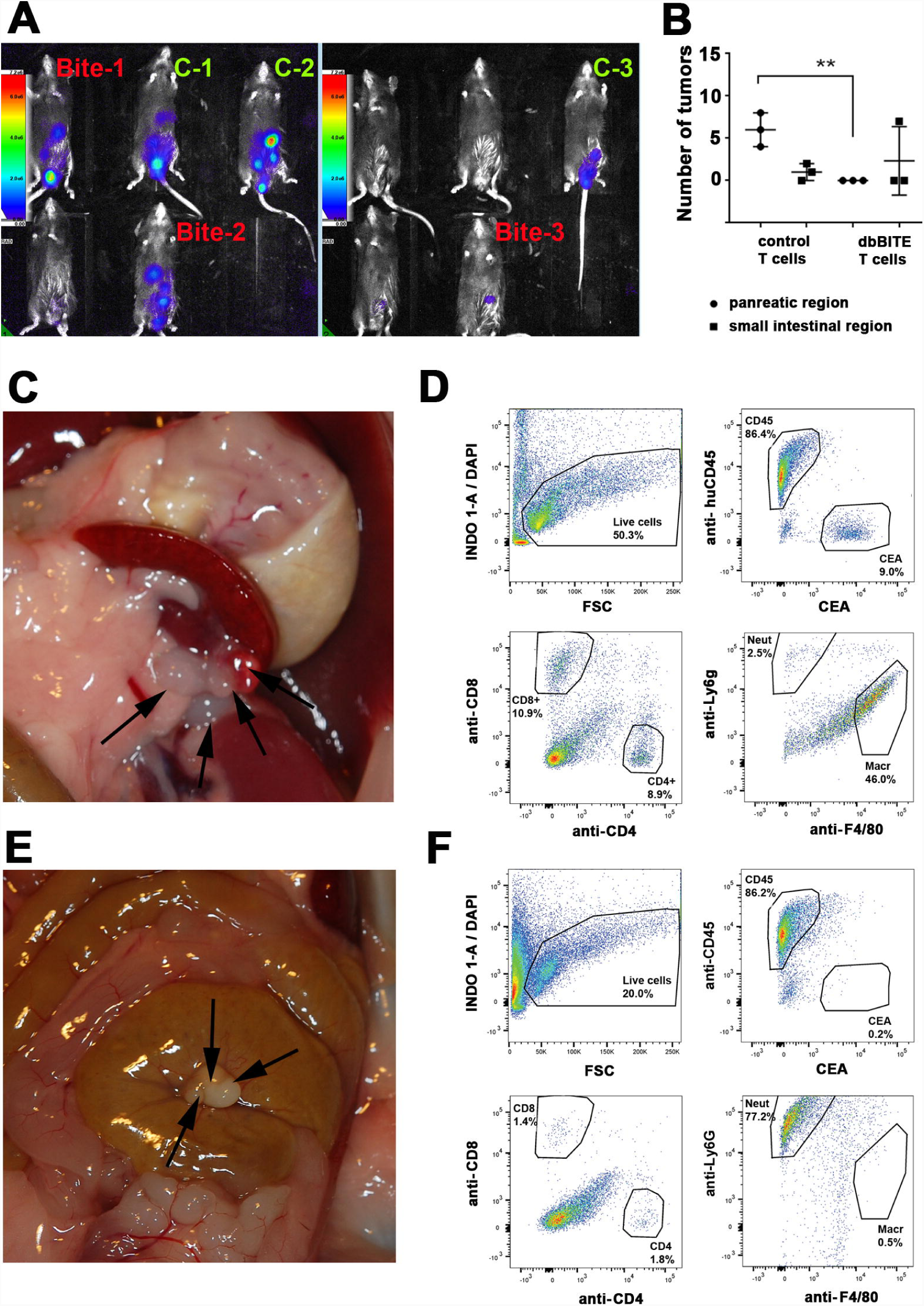
In vivo activity of dbBiTE coated murine T-cells against CEA targets in a CEA transgenic mouse model. **A**. Optical imaging of CEA-Tg mice injected i.p. with mouse carcinoma cell line MC38CEA-Luc at day 9 post injection. Labels: Controls in green and Bite in red indicate group assignment. **B**. Number of tumor nodules in pancreas or intestine region in CEA-Tg mice treated 4x every 3 days with 10 million activated CD3 T cells with or without dbBiTE coating. **C**. Tumor nodules indicated by arrows found in peri-pancreatic location of mouse treated with control T cells. **D**. Flow analysis of digested tumor nodules from animals treated with control T-cells (shown in **C**). Samples were analyzed for presence of DAPI negative live cells, CD45 positive or CEA positive cells. CD45 cells were further analyzed for presence of CD8 and CD4 markers, and CD11b positive myeloid cells were checked for Ly6G and F4/80 markers. **E**. Tumor nodules in peri-intestine location (arrows) of mouse treated with dbBiTE coated T cells. **F**. Flow analysis of tumor nodules (shown in **E**) using same staining and gating as in panel **D**.

## Discussion

This study was initiated to extend the scope of BiTEs beyond the technology requiring genetically engineered dual scFvs, and beyond the approach of coating activated T-cells with a heterogenous mixture of bispecific antibodies. While genetically engineered BiTEs from dual scFv fragments to CD3 and either EpCAM or CD19 have shown significant anti-tumor activity in the clinic [8-10], it is important to extend this approach to readily available intact tumor specific antibodies. In this regard, CEA is an excellent target for many solid tumors including colon, breast, pancreas, and medullary thyroid carcinoma [11]. Although several groups have generated anti-CD3/anti-CEA dual scFv BiTEs [12, 13], their clinically utility is subject to several challenges common to all dual scFv BiTES. A major challenge is that the production of dual scFv BiTEs requires proper protein folding for subsequent large-scale production for clinical use. A second challenge is the need to continuously infuse these BiTEs for therapy due to their rapid blood clearance via the kidney. Since many humanized anti-tumor antibodies have become FDA approved therapeutics, it is highly desirable to generate bispecific antibodies directly from intact antibodies circumventing these two problems. First, there is no need to reengineer the bispecific BiTE, and if an intact anti-tumor IgG is FDA approved, one may obtain it from the pharmacy. Similarly, a murine anti-human CD3 antibody is commercially available and clinically tested [14]. Second, once combined into a bispecific format, their combined molecular weights (300 kda) would preclude blood clearance by the kidney. Although an attractive idea, many attempts to produce such antibodies retaining their dual specifities has been met with difficulty. Among the approaches, knobs-into-holes [15], DuoBody [16], and Cross-Mabs [17], all require re-engineering the parent antibodies. Several chemical cross-linking strategies have been explored, one of which reached the stage of clinical trials. The strategy involved the introduction of protein surface thiols by reaction of surface lysines on IgGs with Traut’s reagent, followed by the hetero-bispecific cross-linking agent sulfosuccinimidyl 4-(*N*-maleimidomethyl) cyclohexane-1-carboxylate [3]. Although the cross-linked product had significant anti-tumor activity when coated on activated T-cells, the product itself was very heterogeneous. A better strategy at producing a more homogeneous cross-linked product was described by Patterson et al. for two IgGs [18] and their Fab fragments [18]. In this approach chemically reduced hinge region cysteines were derivatized with Click reagents that allowed a more specific crosslinking of two different IgGs together. In their study, two clinically available anti-tumor antibodies, anti-EGFR (Cetuximab) and anti-Her2 (Trastuzuab), that were conjugated by cysteine specific Click chemistry retained the specifities of both intact antibodies.

In our study, we demonstrate that cross-linking of murine anti-human OKT3, an IgG2a, to a humanized anti-CEA IgG1 resulted in a bispecific antibody that was isolated in high yield and retained high binding to cells expressing human CD3 or CEA. Importantly, since both antibodies retain both antigen combining arms, they retain the property of avidity, inherent to all intact antibodies, a feature that is lost with dual scFv BiTEs. To emphasize this feature, we have named them dbBiTES for their dual specific, bivalent properties. Since biological products are expected to meet high standards in terms of molecular characterization, it was important to analyze the physical characetictics of dbBiTEs. EM characterization of the resultant 300 kDa particles revealed a flexible six-lobed structure with considerable density at the hinge region, consistent with our expected model. Indeed, the structural model generated using molecular docking and dynamics of the two IgG antibodies was consistent with the EM analysis. Finally, dbBiTE coated human or murine T-cells demonstrated significant and specific anti-CEA cytoxicity for CEA positive target cells. Thus, this approach may be exploited to generate antibody directed activated T cells without the need for genetic engineering, or alternatively, may be used to coat CAR-T cells to give them additional properties.

Since dbBiTEs are at an early stage of development, it’s a task to demonstrate all of their advantages or to predict all of their disadvantages. The presence of intact Fc domains may be viewed as either an advantage or a disadvantage. One clue to their in vivo function was the difference in recruitment of myeloid cells to tumors treated in vivo. dbBiTE coated T-cells appeared to dramatically increase neutrophil infiltration, while in controls, macrophage infiltration dominated. Clearly, more model systems need to be tested to determine if this dichotomy can be generalized and if it truly reflects the Fc activity of dbBiTEs. Overall, we believe this powerful approach can be generalized, since it was demonstrated for both all-human and all-murine systems.

## Acknowledgements

The authors thank the City of Hope Comprehensive Cancer Center (NCI grant P30 CA033572) for support. We acknowledge preliminary EM studies by Marsha Miller and Zhuo Li of the City of Hope EM Core and mass spectrometry by Roger Moore of the City of Hope proteomics core.

## Author contributions

JES conceived the idea. MK, LL, and SB designed and carried out experiments. W-HL, LO, and HL performed in vitro and in vivo studies. Radiolabeling, imaging, and biodistribution studies were performed by JC, KP, NB, and PW. VN supervised and critiqued the molecular dynamics studies. PY provided the humanized anti-CEA antibody M5A and critical review of the ms.

## Materials and methods

### Materials

Dibenzocyclooctyne-amine (BP-22066) and bromoacetamido-PEG_5_-azide (BP-21801) were purchased from Broadpharm (San Diego, CA). Murine anti-human CD3 (OKT3; InVivoMAb, BE0001-2) was from BioXCell (Lebanon, NH), and rat anti-murine CD3 (Cat no 100331, LEAF purified) from Biolegend (San Diego, CA).

### Synthesis of bromoacetamido-DBCO

To dibenzocyclooctyne-amine (DBCO-NH_2_) (27.52mg, 0.1mmol) in 400μl dry DMF was added bromoacetic anhydride (41.0mg, 0.15mmol) and NaHCO_3_ (25.2mg, 0.3mmol). After rotating at RT for 2h under Argon, the mixture was purified by a Gemini C18 column (Phenomenex, CA. 4.6×250mm) on the Agilent (1260 infinity) HPLC instrument using a mobile phase consisting of 0.1% TFA/H2O (solvent A) and 0.1% TFA/Acetonitrile (solvent B), and a linear gradient from 0% B to 95% B in 32min at a flow rate of 1mL/min. The product (18.7mg; yield: 47.1%) was pooled and stored dried at 4°C. The mass of the product was confirmed by ESI mass spectrometry on a Thermo, LTQ-FT; calculated (M+H^+^): 397.27, found: 397.06.

### Reduction of hinge cystines in a IgG antibody

OKT-3 (2mg, 13.3 nmol) in 234μl of PBS was reduced with a 30 molar excess of tris (2-carboxyethyl) phosphine (TCEP) at 37°C for 2h under Argon. The TCEP was removed by desalting on a spin column (Zeba, 7 KDa MW cutoff, Thermo Scientific).

### Alkylation of a reduced antibody with bromoacetamido-DBCO

The reduced OKT-3 was reacted with 20 fold molar excess bromoacetamido-DBCO at RT overnight under Argon. The excess bromoacetamido-DBCO was removed by dialyzing vs PBS (2L x5). The conjugation was confirmed by Agilent 6520 QTOF mass spectrometry. There was one DBCO per light chain and four DBCOs per heavy chain.

### Alkylation of a reduced antibody with bromoacteamido-PEG_5_-azide

Humanized anti-CEA antibody M5A (2mg, 13.33nmol) in 400μl of PBS was reduced with a 30 molar excess of tris (2-carboxyethyl) phosphine (TCEP) at 37°C for 2h under Argon. The TCEP was removed by desalting on a spin column (Zeba, 7KDa MW cutoff, Thermo Scientific). The reduced M5A was reacted with 100 fold molar excess bromoacetamide-PEG_5_-N_3_ at RT overnight under Argon. The excess bromoacetamide-PEG_5_-N_3_ was removed by dialyzing vs PBS (2L x5). The conjugation was confirmed by Agilent 6520 QTOF mass spectrometry. There was one PEG_5_-N_3_ per light chain and 3 per heavy chain.

### Click chemistry and purification by SEC HPLC

OKT-3-DBCO (2mg, 13.33nmol) in 360μl PBS was added M5A-PEG_5_-N_3_ (2mg, 13.3 nmol) in 340μl PBS, pH 7.25. The mixture was rotated at RT for 1h under Argon, then incubated at 4°C overnight. The clicked antibodies were purified by SEC (Superdex 200, 10×300 GL, GE Healthcare) at a flow rate of 0.5ml/min in PBS using a GE AKTAPurifier. Two peaks were collected and concentrated on a 2 mL Vivaspin 10 MWCO (Sartorius, UK). The molecular sizes of peak 1 (300 kDa) and peak 2 (150 kDa) were determined by SDS gel electrophoresis (NuPAGE 4-12% Bis-Tris Gel; Life Technologies, CA) under non-reducing conditions along with antibody standards.

### Transmission electron microscopy and image analysis

dbBiTEs were submitted for analysis to Nanoimaging Services, Inc. (San Diego, CA) Samples were prepared on a thin layer of continuous carbon placed over a C-flat holey carbon grid (2.0/1.0; Protochips). A 3μl drop of purified protein (15 μg/ml in 50 mM Tris/150 mM NaCl, pH 7.5) was applied to a freshly plasma-cleaned grid for 20 seconds, blotted to a thin film using filter paper, and immediately stained with 2% (w/v) uranyl formate for 1 minute. The stain was then blotted away with filter paper and the grid subsequently air-dried. Transmission electron microscopy was performed using an FEI Tecnai T12 electron microscope operating at 120 kV equipped with an FEI Eagle 4k × 4k CCD camera. Images were collected at nominal magnifications of 110,000x (0.1 nm/pixel) and 67,000x (0.16 nm/pixel) using the automated image acquisition software package Leginon [19]. Images were acquired at a nominal underfocus of −1.6 μm to −0.9 μm or −1.4 μm to −0.6 μm and electron doses of approximately 25e-/Å^2^. An example of a typical field at a magnification of 67,000 is shown in **Fig 2A**.

Image processing was performed using the Appion software package [20]. Contrast transfer functions (CTF) of the images were corrected using CTFFind4 [21]. Individual particles in the 67,000x images were selected using automated picking protocols, followed by several rounds of reference-free 2D alignment and classification based on the XMIPP [22] processing package to sort them into self-similar groups. An example of a 2D reconstructed image in shown in **Fig 2B**. A complete set is available upon request.

A random conical tilt 3D structure analysis was performed on 4000 average particles. Over 50 different RCT reconstructions were computed in an attempt to resolve interpretable structures. Two of the best three-dimensional reconstructions of the cross-linked IgG complex resulted in maps at 45-47 Å resolution using between 644-839 particles. The 3D maps of the particles are 150 Å across and reveal structures with five branches of density approximately the size and general shape of Ig domains that define a plane around the periphery of the particle (**Fig 2C**). The position of the sixth Ig domain is disordered in both reconstructions, and is likely either above or below the plane defined by the other Ig domains. The view perpendicular to the plane is flattened in the reconstructed volumes due to limits of the RCT technique, however the central section of the volume in **Figure 2C** had enough additional density to fit most of an Ig domain to sit roughly perpendicular to the orientation of the others, supporting this as the most probable position of the sixth Ig domain in each structure. Density for the Ig domains in each structure varies in both strength and quality and is not sufficient to resolve the hole between the constant and variable domains, and often doesn’t allow for the full size of an Ig domain. This observation, along with blurring in the class averages, is an indication of structural heterogeneity and/or flexibility in the particles, and was a complicating factor in computing an interpretable reconstruction. Due to this limitation, it is not possible to identify Fab vs Fc domains, or determine their connectivity in the reconstructed volumes. A complete set of 3D images is available upon request.

Individual particles were selected using automated picking protocols [20] on both the untilted and tilted images taken at 67,000x. Auto alignment was used to match particles across the tilted image pairs [23]. A reference-free alignment strategy based on the XMIPP processing package [22] was used to separate these particles into classes. RCT geometry [24] was used to reconstruct the 3-D structure of the particle pairs in exemplary class averages. The RCT maps presented in this report are based on 644-839 particles. The nominal resolution of both maps is on the order of ∼45 Å according to the FSC0.5 resolution criterion. The Chimera visualization package [25] was used to produce the surface renderings of the maps.

### Prediction of dbBiTE structure using multiscale simulations

In brief, the two antibody structures were separately modeled after the available crystal structure of mouse immunoglobulin. The docking between the two antibodies was accomplished using coarse-grain MD simulations, imposing attractive forces among the conjugating cysteine residues in the hinge regions of the two antibodies to facilitate binding. The resulting complex structure was then converted to atomistic resolution for comparison with the EM data.

The structures of the M5A and OKT3 antibodies (**Supplementary Fig S2A**) were predicted using the crystal structure of mouse IgG2a as template (PDB ID: 1IGT) [26], using the software MODELLER [27]. For docking, the two antibodies were initially separated by 100Å and positioned with their hinge regions facing each other, as shown in **Supplementary Fig S2A**. The system was then converted to the Martini coarse-grained model [28] (**Supplementary Fig S2B-C**) using the CHARMM-GUI [29]. During the course of molecular dynamics, the internal structures of the Fc and Fab regions were preserved using an elastic network [30], while the linker regions were made completely flexible. This allowed us to model the deformation of the hinge regions as the two antibodies approached each other to form the complex. To facilitate docking, we imposed distance restraints between the cysteine residues in the hinge regions of the two antibodies. Each M5A monomer has two cysteines in the lower hinge region connecting heavy chains, while OKT3 has three cysteines connecting heavy chains. We restrained the Cα carbons of the hinge region cysteines of the two antibodies using a flat-bottomed potential with an equilibrium distance of 15Å. Beyond 15Å, the restrained residues experienced an attractive elastic force with a force constant of 1000 KJ/mol, while no forces were experienced below 15Å. The equilibrium distance for the restraint between the conjugated cysteines was determined from MD simulations of two cysteine residues connected by the click chemistry linker in aqueous solution. The equilibrium distance was chosen as the mean value in the distance distribution of the Cα carbons in this system (**Supplementary Fig S2D**). We hypothesized that upon docking, at least one cysteine in the hinge region of one antibody will be conjugated to one cysteine of the other antibody via click chemistry. To recreate this condition, the system was simulated imposing the criterion that at least one of the distance restraints between the cysteine residues should be satisfied during dynamics.

The system was minimized for 6000 steps using steepest descent minimization, then heated up to 303K, followed by equilibration in the NPT ensemble for 10 ns (303K, 1 atm). During equilibration, the protein atoms were restrained at their initial positions with a force constant of 4000 KJ/mol. The production MD was performed at 303K in the NPT ensemble for 500 ns using a time step of 20 ps, by imposing the distance restraints as described above. The MD simulations were performed using GROMACS 5.1 [31]. The final structure from the coarse-grained MD simulation (**Supplementary Fig S2E**) was converted to the all-atom structure using the scheme described in ref. [32] and visualized using PyMOL [33].

### Tumor cell lines culture and T cell activation

Human breast carcinoma MDA-MB-231 and murine breast carcinoma cell line E0771 and colon cancer cell line MC38 were stably transfected with a CEA expressing plasmid as described earlier [7]. All cell lines were maintained in DMEM supplemented with 10% fetal bovine serum and 100 U/ml penicillin/streptomycin.

Human PBMC were isolated by centrifugation of whole blood on Ficol-Paque (GE Healthcare) gradient for 30 minutes (500g) and washed with PBS 3 times. After counting and checking cells viability PBMCs were plated on 6 well plate, previously coated with anti-CD3 antibody (2ug/mL for 2h at 37°C, washed once with PBS at concentration 2×10^6/mL in RPMI1640 containing 10% FBS and 100U/ml of recombinant human IL-2 (BioLegend, San Diego, CA). After 72h, cultures of activated cells were expanded by platting 5×10^5/ml in RPMI1640 containing 10% FBS and 100U/ml of recombinant human IL-2. Cells were cultured for up to 7 days and used for functional experiments *in vitro*.

Mouse CD3 positive T cells were isolated from spleens collected from CEA transgenic mice using negative selection kit (Stemcell Technologies, Canada) and plated for activation on 6 well plates previously coated with αCD3 antibody (BioLegend, CA; 2ug/ml for 2h in 37C, washed once with PBS), at concentration 2×10^6/ml in IMDM containing 10% FBS and 100U/ml of recombinant mouse IL-2 (BioLegend, CA). Cells were cultured for up to 7 days and used for functional experiments *in vitro* and *in vivo*.

### Coating conditions for dbBiTEs onto activated human or mouse T cells

Both target cells (human tumor cell line: MDA-MB-231±CEA, mouse tumor cell lines: E0771±CEA, MC38±CEA) and T cells (human PBMCs derived T cells and mouse CD3 positive cells) were first incubated in 1% goat serum blocking solution, then stained on ice with 1ug/mL of dbBiTE in PBS containing 1% goat serum for 30 minutes. After washing with PBS, cells were stained on ice for 30 minutes with 2ug/mL of secondary antibodies (all from ThermoFisher): goat anti-mouse Alexa555 and goat anti-human Alexa647 for OKT3-M5A dbBiTE; goat anti-rat Alexa555 and goat anti-human Alexa647 antibodies for mCD3-M5A dbBiTE. After washing, cells were resuspended in PBS and analyzed by FACS (LSRFortesa X-20, BD **Biosciences). Unstained cells,** or cells stained with **secondary** antibodies only served as negative controls. OKT3 and rat anti mCD3 (BioLegend, CA) and M5A were used as positive controls followed by secondary antibodies staining performed as described above.

### In vitro cytoxicity assay

Target cells were plated on 96-well plates at concentration 10×10^3 per well in 100uL of DMEM containing 10% FBS. After 3 or 18 hours, depending on cell line, 100uL of human activated and antibody coated PBMCs or mouse activated and antibody coated CD3 T cells were added to target cells in concentrations corresponding to final effector to target ratios 10:1, 5:1, 2.5:1 and 1.25:1. Controls with uncoated or anti-CD3 coated cells were used in the same ratios. Cells were co-incubated for 18 hours at 37°C followed by analysis of lactate dehydrogenase (LDH) release. The assay was performed following manufacturer’s protocol (Takara, Japan) and relative cytotoxicity was calculated according to the same protocol.

### *In vivo* tumor targeting with dbBiTE coated activated mouse T cells

These animal experiments were performed using CEA transgenic mice generated in Beckman Research Institute at City of Hope National Medical Center as described before [7]. Mouse care and experimental procedures were performed under pathogen-free conditions in accordance with established institutional guidance and approved protocols from the Institutional Animal Care and Use Committee of Beckman Research Institute at City of Hope National Medical Center. For the colon cancer model, MC38/CEA/Luc cells were injected subcutaneously at the concentration of 2×10^6^. At day 9 post injection optical imaging was performed using Lago imaging system (Spectral Instruments Imaging, AZ) and treatment groups were assigned. Mice were treated 4x every 3 days with 10 million activated CD3 T cells with or without dbBITE coating. Four days after last T cells injection the tumor nodules were counted and removed, then dissociated by enzymatic digestion using gentleMacs Octo Dissociator and dissociation kit following the manufacture’s protocol (Miltenyi Biotec). Samples were analyzed by flow for presence of DAPI negative, live cells, CD45 positive or CEA positive cells. CD45 cells were further analyzed for CD8 and CD4 markers, and CD11b positive myeloid cells were analyzed for Ly6G (neutrophil) and F4/80 (macrophage) markers.

### PET imaging of a CEA positive tumor with ^64^Cu radiolabed DOTA-dbBiTE

dbBiTEs were conjugated with NHS-DOTA as previously described [6]. Immunoreactivity to CEA was confirmed by addition of ^64^Cu-DOTA-dbBiTE to CEA followed by SEC as shown in **supplementary Fig S5C**. Animal imaging studies were performed in NOD/SCID mice bearing s.c. LS-174T CEA positive tumors as previously described [6]. PET scans were acquired with an Inveon microPET/CT scanner (Siemens Medical Solutions). Mice were anesthetized with 2%–4% isoflurane in oxygen, placed on the PET scanner, and injected with a single intravenous dose of 100 μCi (10 μg) of ^64^Cu-DOTA-dbBiTE in 1% human serum albumin–buffered saline through a tail vein catheter. At the terminal time point, the mice were euthanized and biodistribution studies performed on the tissues indicated in **supplementary Fig S6**.

