## Supplemental Figures for "Generation of dual specific bivalent BiTEs (dbBIspecific T-cell Engaging antibodies) for cellular immunotherapy"

### SUPPLEMENTAL MATERIAL

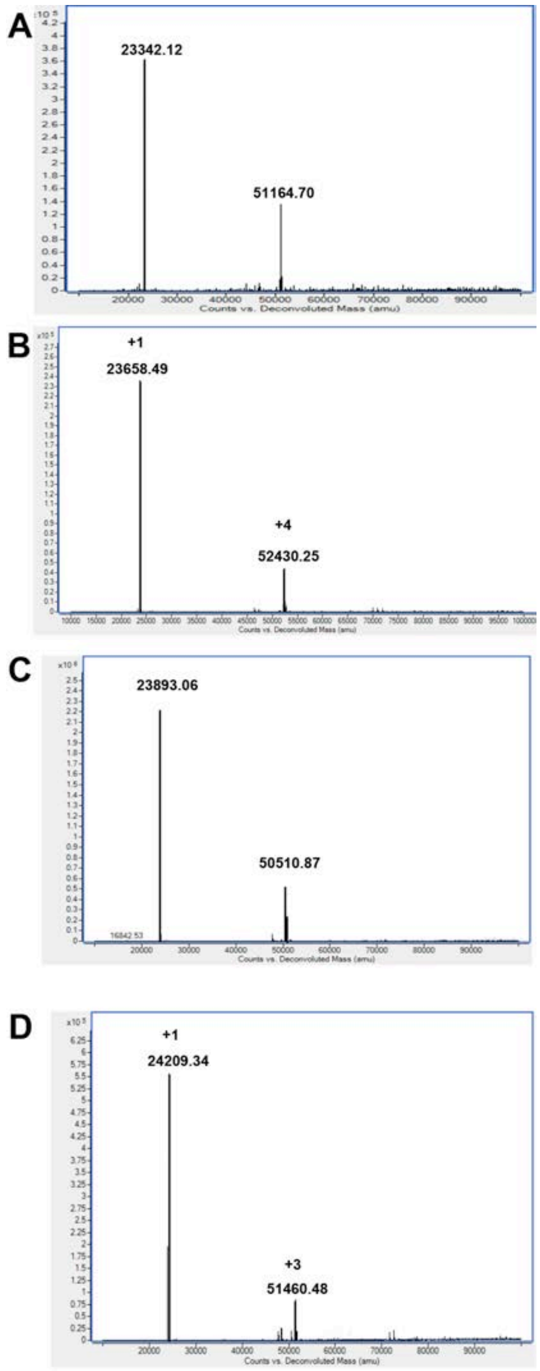


**Supplemental Figure S1. ESI MS analysis of DBCO and $PEG_5$ azido derivatized antibodies**. **A-B**. Anti-CD3 antibody OKT3 before (**A**) and after (**B**) derivatization with a bromoacetamido DBCO. **C-D**. Anti-CEA antibody M5A before (**C**) and after (**D**) derivatization with bromoacetamido-$PEG_5$-azide.

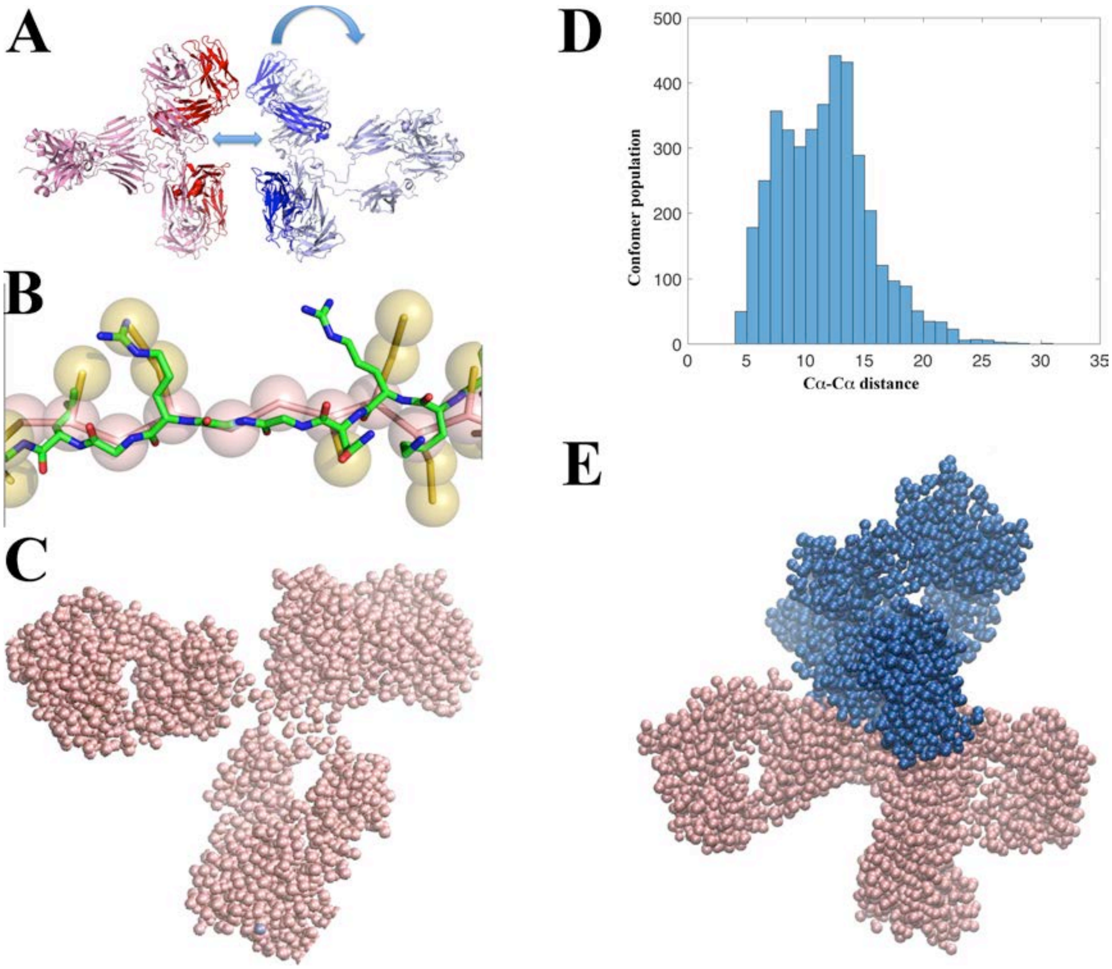


**Supplemental Figure S2. Molecular dynamic simulation of the formation of dbBiTES**. **A**. The x-ray structures of two IgG1 antibodies were aligned and one rotated 90$^o$ to allow them to approach each other. **B**. Each antibody was converted to a coarse grain model in which each amino acid is converted to a sphere to simplify computational analysis. A section of one antibody is shown to indicated the principle. **C**. A complete coarse grain model of a single IgG1. **D**. Alpha carbon distances (in angstroms) in the hinge region as the two coarse grain models are docked. **E**. The final model in which at least two cysteines in the hinge regions have been docked.

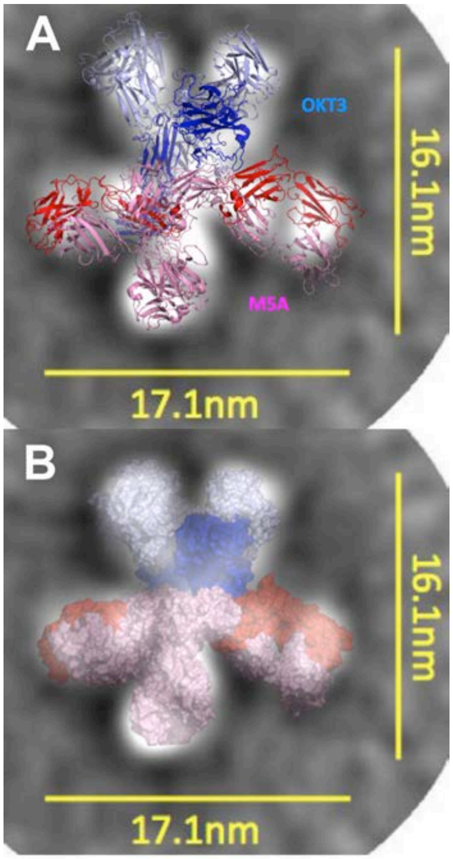


**Supplemental Figure S3. Superposition of dbBiTE model onto an average EM particle**. **A**. Atomic structure constrained to average EM particle. **B**. Space filling model of a dbBiTE constrained to an average EM particle.

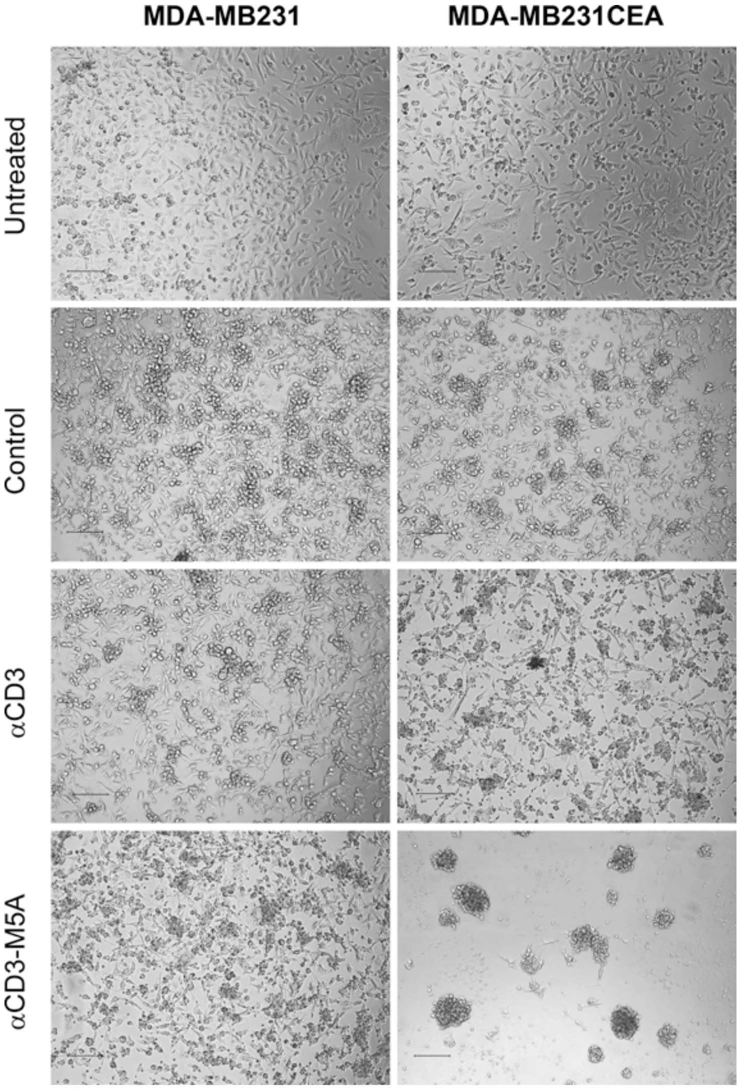


**Supplementary Fig S4. Microscopic imaging of dbBiTE coated T-cells killing target cells**. MDA-MB231 ±CEA cells were treated with dbBiTE coated activated human T-cells at an E:T ratio of 10:1 for 24 hrs and visualized by microscopy. Activated T-cells coated with anti-CD3 alone caused some clumping and killing of targets, but dbBiTE coated T-cells completely killed all of the CEA+ target cells compared to the CEA- target cells.

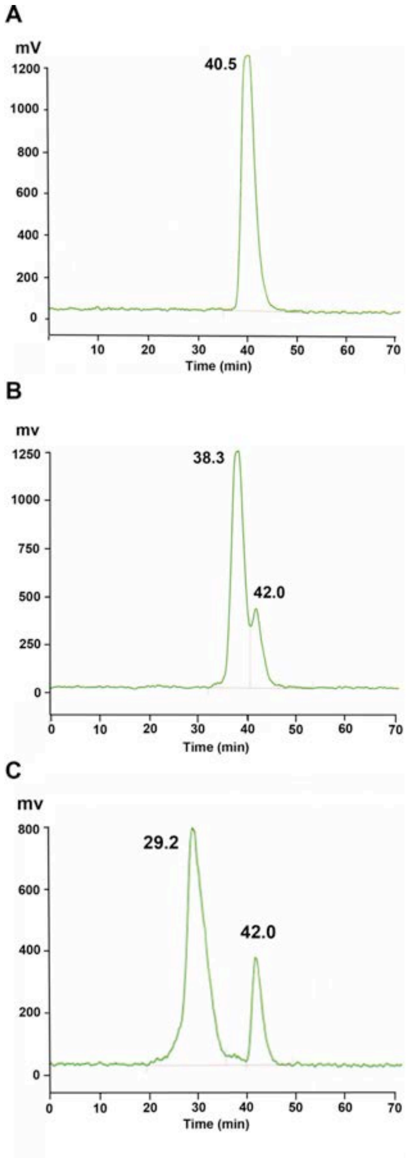


**Supplementary Fig S5. SEC analysis of $^{64}$Cu-DOTA labeled M5A and dbBiTE. A.** Analysis of radiolabeled M5A by SEC. **B**. Analysis of radiolabeled dbBiTE by SEC. **C**. Analysis of radiolabeled dbBiTE after the addition of a 20 fold excess of CEA. Radiolabeled dbBiTE (300 kDa) contains about 23% of 150 kDa material, or 11.5% on a molar basis.

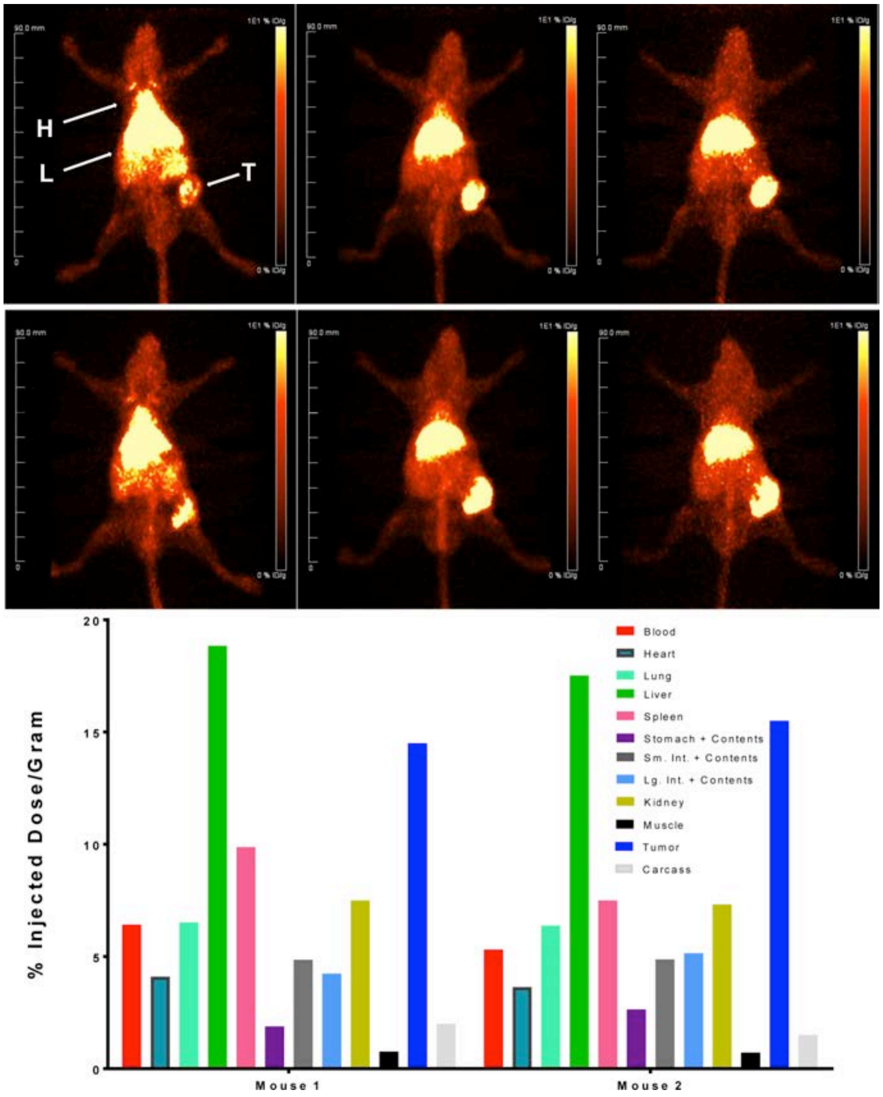


**Supplemental Figure S6. In vivo targeting of $^{64}$Cu-DOTA-dbBiTE in NOD-SCID mice bearing CEA positive LS-174T tumors**. Upper two panels show two mice imaged at 4, 20, and 44hr. Organs labeled are heart (H), liver (L) and tumor (T). Lower panel shows biodistribution of indicated tissues at terminal imaging time point.

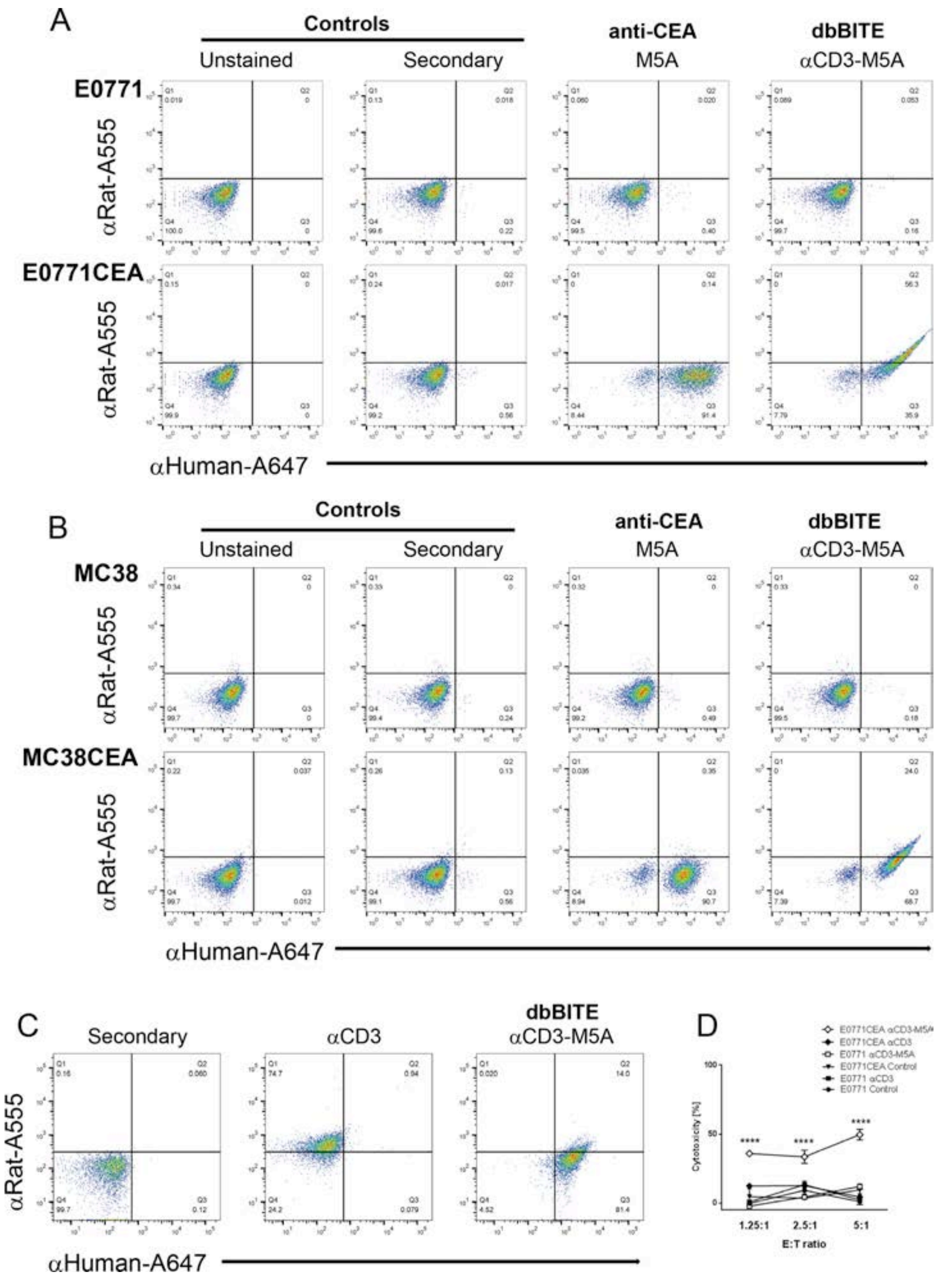


**Supplementary Fig S7. Binding of dbBiTE rat-anti-murine CD3-human anti-CEA (M5A) to murine target cells ±CEA and cytotoxicity. A**. Murine breast cancer cell line E0771 with or without transfection with CEA were incubated with M5A or dbBiTE and stained with secondary antibodies specific for rat or human IgG. Controls show lack of binding of secondary antibodies. M5A binds only to CEA+ target with no binding of the rat secondary antibody, while dbBiTE coated targets bind both secondary antibodies. **B**. Analogous experiments with murine colon carcinoma cell line MC38 with or without transfected CEA. **C**. anti-CD3 coated activated murine T-cells bind rat secondary antibody only, while dbBiTE coated abtivated murine T-cells bind both secondary antibodies. **D**. dbBiTE coated T-cells exhibit significant cytotoxicity to CEA+ targets vs CEA- controls.

**Supplemental movie SM1**. **Coarse grain simulation of docking of two antibodies.** OKT3 is shown in blue, while M5A is shown in red; the spheres represent coarse grain collections of protein atoms as explained in **supplementary Figure S2B**.

**Supplemental movie SM2. Predicted atomistic structure of dbBITE using multiscale simulations.** OKT3 is shown in blue, while M5A is shown in red; the click chemistry linkers connecting the hinge region cystines in both antibodies are shown as green sticks; the cystine residues are shown as spheres.

**Supplemental movie SM3. Predicted atomistic structure of dbBITE shown in molecular surface representation.** The color scheme is as follows; OKT3: light chain – blue, heavy chain – cyan; M5A: light chain – red, heavy chain – yellow.

**Supplemental movie SM4**. **Time lapse photography of dbBiTE coated human T-cells (1 µg/10M cells) killing MDA MB231/CEA target cells.** (E:T of 10:1) over 18 hrs.

**Supplemental movie SM5**. **Time lapse photography of dbBiTE coated human T-cells (1 µg/10M cells) incubated with MDA MB231 control cells.** (E:T of 10:1) over 18 hrs.
